# The tropical coral *Pocillopora acuta* has a mosaic DNA methylome, an unusual chromatin structure and shows histone H3 clipping

**DOI:** 10.1101/722322

**Authors:** David Roquis, Ariadna Picart Picolo, Kelly Brener Raffalli, Pascal Romans, Patrick Masanet, Céline Cosseau, Guillaume Mitta, Christoph Grunau, Jeremie Vidal-Dupiol

**Affiliations:** Université de Perpignan Via Domitia, CNRS, Ifremer, Université de Montpellier, Perpignan, F-66860, France; CNRS-UPMC UMS 2348, Observatoire Océanologique de Banyuls, Université Pierre et Marie Curie - Paris 6, France; Aquarium de Canet-en-Roussillon SPL Sillage, F-66140 Canet-en-Roussillon, France

**Keywords:** *Pocillopora acuta*, *Pocillopora damicornis*, coral epigenetic, chromatin structure, mosaic DNA methylation, Histone H3 clipping

## Abstract

*Pocillopora acuta* is a hermatypic coral with a worldwide distribution and a strong ecological importance. Anthropogenic disturbances and global warming threaten it. Thermal stress can induce coral bleaching, a phenomenon in which the mutualistic symbiosis between the coral polyps host and its endosymbiotic unicellular algae is disrupted, and can lead to the death of entire colonies. Previous works have shown that soma clonal colonies display different levels of survival depending on the environmental conditions they previously faced. Epigenetic mechanisms are good candidates to explain this phenomenon. The clonal nature of a colony and the possibility of generating genetically identical colonies through propagation make corals an attractive model to study the impact of the environment on the epigenome. However, until now, no work had been published on the *P. acuta* epigenome. One of the main problems is caused by the intracellular location of Symbiodinium, which makes it complicated to isolate coral chromatin free of contamination by endiosymbiotic biological material. Here, (i) we describe a simple method to purify *P. acuta* chromatin, (ii) we provide the first description of a coral methylome, with a mosaic pattern of cytosine methylation principally in a CpG context (4% of all CpG), and (iii) we show that *P. acuta*, but not all corals, has an unusual chromatin structure, and displays histone H3 clipping.

## Introduction

Epigenetic modifications are good candidates to explain rapid, inheritable and reversible phenotypes without change in the DNA sequence (Danchin et al., 2011). They range from chemical modifications of DNA (eg. cytosine methylation), covalent changes of proteins participating to chromatin structure (e.g. histone modifications), as well as nuclear localization of chromosomes, and short untranslated RNA involved in post-transcriptional silencing of genes and repeated regions (Wu and Morris, 2001). These modifications have an impact on the chromatin structure leading to the modulation of transcriptional activity. It is now clear that environmental factors can influence the epigenome to induce the expression of new phenotypes (Duncan et al., 2014; Feil and Fraga, 2012). In natural populations with genetic diversity, disentangling the precise respective roles of genetics and epigenetics in environmentally triggered phenotypes is hardly feasible, if not impossible. Despite the ecological importance of corals, technical challenges have until now prevented corals to serve as model species to study the effect of environment on epigenome. Indeed, although several authors proposes to investigate epigenetic mechanisms possibly related to adaptation to climate change in hermatypic corals (Palumbi et al., 2014; Putnam et al., 2016; van Oppen et al., 2015), literature is scarce, and we found only few publications focusing on this matter (e.g. Dimond et al. 2015, 2016, 2017). In addition to the absence of reference genome, many technical challenges make the epigenome analysis difficult: extraction of cells and tissues from the stony exoskeleton, working at a marine salt concentration to maintain cell and chromatin integrity, and most importantly, separation of coral and *Symbiodinium* biological material. *Symbiodinium* are endocellular, and to focus on the coral epigenetic modifications, we can focus on non-symbiotic life stages, use naturally or artificially bleached colonies, or use laboratory techniques enabling physical separation of symbionts and host’s nuclei and genetic material.

In this article, we describe a method we developed to isolate coral nuclei of *Pocillopora acuta*, and provide a first insight about the chromatin structure of this species. We chose this coral for its worldwide distribution and ecological importance (IUCN, 2015), its fast growth rate allowing to quickly generate new colonies through propagations (Hughes et al., 2015), and its documented response to thermal stresses (Vidal-Dupiol et al., 2009; Vidal-Dupiol et al., 2014). *P. acuta* has a genome size of approximately 325 Mb (Vidal-Dupiol et al. in preparation), while *Symbiodinium* genome size is estimated at ~1,200-1,500 Mb (Lin et al., 2015; Shoguchi et al., 2013), which means that a single *Symbiodinium* brings as much DNA as five coral cells, hence the necessity to reduce contamination from endosymbiontic material at the lowest possible level. We used then the isolated coral nuclei for the analysis of epigenetic information carriers. We showed that mosaic cytosine methylation occurs and that 5-methyl-cytosine is predominantly found in CpGs. We also found that canonical histone modifications exist, and that *P. acuta* has an unusual chromatin structure.

## Material & Methods

### Biological material

The *Pocillopora acuta* isolate used in the present study was harvested in Lombock (Indonesia, CITES number: 06832/VI/SATS/LN/2001) and maintained at the Banyuls Aquarium (France) under optimal conditions. Previously assigned to *Pocillopora damicornis* this isolate was reassigned to *P. acuta* (Vidal-Dupiol et al. Supplementary article 1). This assignation was based on the 840 based pair sequence of the ORF marker that enable to separate *P. acuta* and *P. damicornis* (Schmidt-Roach et al., 2014). Bleached coral colonies were obtained through a menthol treatment. Briefly, the colonies were placed in a four-liter tank filled with seawater. Water motion was created using a submerged water pump (100 L/h), temperature was maintained at 27°C and light adjusted to 75 μmol/m²/s (PAR). The protocol for the menthol treatment was adapted from (Wang et al., 2012), the first day, the corals were subjected to a concentration of 0.58 mmol/L for 6h. After this exposure, they were transferred to the coral nursery for a 18h recovery period. During the second day, the same protocol was applied (menthol treatment and recovery step). The third day, the corals were exposed again to the same treatment but only until the polyps were closed. Once the polyps were closed the corals were placed in the coral nursery for recovery and zooxanthellae loose. This last step typically takes four to five days. Once the polyp open up again samples were taken. Healthy and bleached coral fragments were immediately frozen and stored in liquid nitrogen.

In this work, we also employed other *Cnidaria* for comparison in the chromatin extraction and digestion experiments. We chose three other hermatypic corals, *Stylophora pistillata* (CITES number IAZ3924), *Montipora digitata* (CITES number IEZ0069) and *Euphyllia divisa* (CITES number IUZ1609). We also used *Aiptasia sp*. and *Anemonia manjano*, two sea anemones also from the *Cnidaria* phylum, sharing an endosymbiodic relationship with *Symbiodinium*, but without an aragonite skeleton. All these species were acquired from the Canet en Roussillon Aquarium (France). All of the species we used in this study are part of the *hexacorallia* subclass. The anemones are from the *actinaria* order, and the corals from the *scleractinia* order.

*Symbiodinium* cultures (clade C1/F) were obtained from the Roscoff Culture Collection (strain CCMP 2246, cat# RCC4017).

### In-Silico identification of canonical histones

We were interested to use isolated coral chromatin to perform chromatin immunoprecipitation (ChIP) on histone modifications. Before starting experiments, we wanted to confirm that histones were present in *P. acuta* genome, and that their amino acid sequence would be similar enough to other metazoan to be able to target them with commercial antibodies. We searched the *P. acuta* transcriptome (Vidal-Dupiol et al., 2014; Vidal-Dupiol et al., 2013) for transcripts annotated as histones H2A, H2B, H3, H4. These sequences were translated to amino-acid sequences and aligned with MEGA 7 (Tamura et al., 2013) to reference protein sequences from *Mus musculus*, *Schistosoma mansoni*, a platyhelminth for which our laboratory has extensive experience in ChIP, and three other *Cnidaria*: *Hydra vulgaris, Nematostella vectensis* and *Acropora digitifera* (scleractinia coral, like *P. acuta*). Accession numbers for each species are in **Table 1**.

**Table 1:** List of accession numbers used to compare *P. acuta* canonical histone sequences

|  | H2A | H2B | H3 | H4 |
| --- | --- | --- | --- | --- |
| *N. vectensis* | XP_001626711.1 | XP_001617661.1 | XP_001623766.1 | EDO36726.1 |
| *H. vulgaris* | XP_002158325.1 | XP_002156085.2 | XP_002154470.1 | XP_004205459.1 |
| *A. digitifera* | XP_015747526.1 | XP_015757821.1 | XP_015766372 | XP_015748868.1 |
| *M. musculus* | AAA37763.1 | NP_783594.1 | NP_038576.1 | NP_291074.1 |
| *S. mansoni* | CCD78657.1 | CAY18308.1 | CAZ35552.1 | CCD74906.1 |

### Western blots

As we intended to perform chromatin immunoprecipitation targeting histone modifications, we first did western blots using commercial anti-histones antibody to assess their efficiency and specificity on *P. acuta* canonical and modified histones. Histones were purified with the abcam histone extraction kit (ab113476) from bleached and healthy coral fragments of approximately 5 mm of diameter (~ 0.5 g). Alternatively, for rapid extraction, coral fragments were put in 1.5 mL tubes filled with 500 μL of a solution containing 62.5 mM TRIS/Cl pH 6.8, 3% SDS, 10% sucrose, 0.2 M dithiotreitol (DTT) and 1.25 mM sodium butyrate. Tubes were sonicated using a Vibra Cell 75185 at 70% amplitude, 3 times 15 seconds, on ice. A cooldown on ice of 30 seconds was done between each sonication. *Dinoflagellata* phylum, which includes *Symbiodinium*, does not possess canonical histones (Rizzo, 2003) and was used as a negative control. We chose hamster (*Mesocricetus auratus*) as a positive control, as most antibodies are commercially developed and tested on mammals. Hamster brain and cultured aposymbiotic *Symbiodinium* protein extracts were processed the same way as *P. acuta*. The laboratory has permission A 66040 from both French Ministère de l’agriculture et de la pêche and French Ministère de l’Education Nationale de la Recherche et de la Technologie for experiments on animals and certificate for animal experimentation (authorization 007083, decree 87-848 and 2012201-0008) for the experimenters. Housing, breeding and animal care follow the national ethical requirements.

The extract was cleared by centrifugation for 30 minutes at 1500 g, and the supernatant was collected. Total proteins (5 mg per sample) were mixed with Laemmli buffer (final concentration 1X) and denatured at 99°C for 5 mins. ECL Full-Range Rainbow (Amersham cat# RPN800E) was used as a molecular weight marker. Protein separation was done on 10% SDS-PAGE gel electrophoresis before being blotted on a nitrocellulose membrane (Trans-Blot turbo, Bio-Rad). The membrane was blocked with 5% non-fat dry milk in TBST (TBS buffer containing 0.05% tween 20) one hour at room temperature. One of the following primary antibody, diluted in 5% non-fat milk in TBST was used for each western blot: anti-histone H3 (Abcam cat# ab1791 lot# 784471, dilution 1/500), anti-H3K36me2 (Abcam cat# ab1220 lot# GR75522, dilution 1/500), and anti-H3K27me3 (Diagenode, cat#C15410069, lot# A1821D). After incubation, membrane was washed 3 times for 10 minutes in TBST. It was incubated with secondary antibody (peroxidase conjugated, goat purified anti-rabbit IgG [Pierce cat# 31460, lot# HB987318]) diluted 1/5000 in TBST for 1 hour. After washing 3 times for 10 minutes in TBST, the detection was carried out using the ECL reagents and the ChemiDoc MP Imaging system (BioRad). Estimation of the relation between number of amino acid residues and molecular weight in kD was done with http://www.calctool.org/CALC/prof/bio/protein_size.

### Coral nuclei isolation

Our chromatin extraction protocol is based on (Cosseau and Grunau, 2011), but had to be adapted so that salinity, ionic strength and osmolarity of various buffers would match those of seawater. *P. acuta* (healthy or bleached), *M. digitata, S. pistillata* and *E. divisa.* polyps were removed from the aragonite skeleton using an Airpick. This instrument is custom-designed to gently blow air through a pipet tip. Advantage of the Airpick is that it can be used with small volume of extraction solution and with a low amount of biological material. Extractions with Airpick were performed on ice for 5 minutes (or until no tissues were left on the skeleton) with the coral fragments (approximately 5 mm of diameter and 0.5 g each) inside a 50 mL tubes or small sealed plastic bags filled with 10 mL of a buffer 1 (See **Table 2** for composition of all the buffers used in this article). Tubes were then centrifuged at 800 g at 4°C for 10 minutes. Supernatant was carefully removed, and pellets were gently resuspended in 1 mL of buffer 1 and 1 mL of buffer 2. The mix was transferred in Dounce homogenizer and ground on ice for 4 minutes with pestle A, and then left to rest on ice for 7 minutes. Liquid was transferred on corex tubes already containing 8 mL of buffer 3, in a manner that the homogenate would form a layer on top of buffer 3. The two solutions have a different density, which allow them to stay one on top of the other without mixing. Corex tubes were centrifuged 20 mins at 4°C and 7,800 g, with the lowest break possible. This step allows separating *P. acuta* nuclei, which will form a pellet at the bottom of the tube, from cell debris and *Symbiodinium*, which stay at the interphase in the homogenate on top of buffer 3. Supernatant was completely removed by pouring out of the tubes and then by aspiration with micropipette. Pellets, containing coral nuclei and chromatin, were either used for DNA purification (to use for bisulfite sequencing) or chromatin digestion (with the objective of using digested chromatin for ChIP).

200 μg of *Aiptasia sp.* and *A. manjano* samples were directly ground in Dounce homogenizer with 1 mL of buffers 1 & 2, and then processed the same way as *P. acuta.*

### Chromatin digestion

One of the most important steps to study histone modifications through chromatin immunoprecipitation is chromatin shearing. This can be done either with crosslinking followed by sonication, or natively using micrococcal nuclease (*M*Nase). Crosslinking creates covalent bounds between DNA and proteins interacting with it, and can be broken in di or tri nucleosomal fragments through timed sonication. This method is usually employed for transcription factor analysis, but is not needed in the case of histones, as they are strongly bound to DNA. Crosslinking also has the downside of making the precipitation very inefficient (partly because it changes the structure of the epitopes and reduce the recognition by antibodies) and can also fix some transient interactions with minor functional significance (Das et al., 2004; Turner, 2001). We favored the native approach with *M*Nase, a bacterial enzyme digesting DNA between nucleosomes. Digestion time has to be optimized to obtain only mono-to penta-nucleosomal fragments. To do so, nuclei pellets (bleached and healthy coral, *A. pallida, A. manjano* and *D. magna*) from step 2.4 were resuspended in 1.5 mL of *M*Nase digestion buffer (see **Table 2** for composition).

**Table 2:**
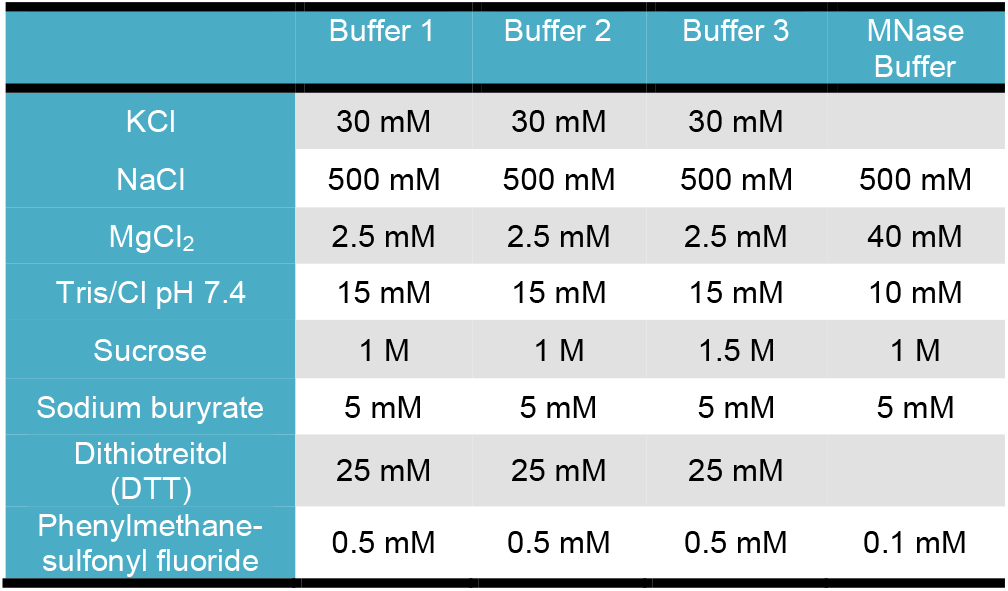
Final concentration for all components of the various buffers used in the chromatin extraction and micrococcal nuclease digestion.

|  | Buffer 1 | Buffer 2 | Buffer 3 | MNase Buffer |
| --- | --- | --- | --- | --- |
| KCl | 30 mM | 30 mM | 30 mM |  |
| NaCl | 500 mM | 500 mM | 500 mM | 500 mM |
| MgCl<sub>2</sub> | 2.5 mM | 2.5 mM | 2.5 mM | 40 mM |
| Tris/Cl pH 7.4 | 15 mM | 15 mM | 15 mM | 10 mM |
| Sucrose | 1 M | 1 M | 1.5 M | 1 M |
| Sodium butyrate | 5 mM | 5 mM | 5 mM | 5 mM |
| Dithiothreitol (DTT) | 25 mM | 25 mM | 25 mM |  |
| Phenylmethane-sulfonyl fluoride | 0.5 mM | 0.5 mM | 0.5 mM | 0.1 mM |

Twenty microliters of the solution were kept for a coral nuclei and *Symbiodinium* count (See step below). Aliquots of 250 μL were prepared in Eppendorf tubes for digestion. 1 μL of 1 U/μL *M*Nase (Affymetrix cat# 70196Y) was added to each tube, and incubation at 37°C was done for 0, 2, 4, 6 and 8 minutes. We also prepared a negative control with no *M*Nase and incubed at 37°C for 8 minutes. Because *Aiptasia* sp., *A. manjano*, *M. digitata*, *E. divisa,* and *S. pistillata*, were used with a comparison purpose, we only performed the digestion with three time points: 0, 4 and 8 minutes. Digestion was stopped with the addition of 20 μL of 1M EDTA and tubes were left on ice for 5 minutes. The soluble fraction of chromatin was separated from cell debris by centrifugation at maximum speed (13,000 g) 10 minutes at 4°C, and DNA in the supernatant was purified using QIAquick PCR Purification kit (Qiagen cat# 28104). DNA fragments were eluted in 30 μL of TE buffer (supplied with the kit). Twenty to 30 μL of the purified DNA fragments were then separated by electrophoresis through a 1.8 % agarose gel stained with ethidium bromide for 60 minutes at 100 V (in 0.5X Tris Borate EDTA buffer).

### Estimation of P. acuta nuclei and intact Symbiodinium in chromatin pellets

We used a chromatin extraction method which was gentle enough to break *P. acuta* cytoplasmic membrane without lysing their nuclei or *Symbiodinium*. Coral nuclei were then isolated from cell debris and *Symbiodinium* due to the density difference of the solutions used during the centrifugation step in corex tubes (step above). To get an estimation of the amount of possible *Symbiodinium* contamination in the nuclei/chromatin pellet when using healthy *P. acuta* as starting material, we took 20 μL of coral nuclei resuspendended in *M*Nase digestion buffer and added Hoechst 33342 (Invitrogen cat# C10329) at a final concentration of 1/2,000. We put 10 μL on a microscope slide and observed on a fluorescent microscope at 350 nm. Hoechst 33342 stains coral nuclei with a blue fluorescence, but does not enter intact *Symbiodiniums* (red fluorescence due to chlorophyll).

### DNA purification and whole genome bisulfite sequencing (WGBS)

Bisulfite sequencing needs DNA to be completely protein-free. To do so, nuclei pellets obtained at step 2.4 were resuspended in 180 μl of the ATL buffer from the DNeasy Blood and Tissues Kit (Qiagen cat# 69504) and then processed as described in the manufacturer documentation. We sent the purified DNA from three healthy coral colonies for bisulfite sequencing (BS-Seq) to GATC biotech (www.gatc-biotech.com), which performed both bisulfite conversion and sequencing. Libraries with fragment size ranging from 150 to 500 bp were paired-end sequenced (100 bp on each extremity) on an Illumina Hi-Seq 2000.

### Quality control, alignment on P. acuta genome and methylome description

All data treatment was carried out under a local galaxy instance (Goecks et al., 2010) (http://bioinfo.univ-perp.fr). Fastq Groomer v1.0 was used for verification of the fastqsanger format, and the FASTX-Toolkit v0.0.13 (Compute quality statistics, Draw quality score boxplot, Draw nucleotides distribution chart) was used for initial quality control. First and last five nucleotides for all reads were removed using the TRIM tool from the FASTX-Toolkit because of lower quality. Overall, read quality was judged sufficiently good (the majority of reads showed a quality score above 26 for the non-trimmed position) and no further quality filter was applied.

Read alignment on the *P. acuta* reference genome (Vidal-Dupiol et al. in preparation) was done in paired-end using Bismark v2.74 (Krueger and Andrews, 2011) and Bowtie2 (Langmead and Salzberg, 2012) with default parameters. We only kept alignments for which both read mates were successfully aligned and fragment size was between 150 and 500 bp. BAM files output from Bismark were sorted and converted to SAM format using Samtools v0.1.19 (Li et al., 2009). SAM files were loaded into the R package methylKit v0.5.6 (Akalin et al., 2012) using the read.bismark() command and specifying the cytosine context (CpG, CHG or CHH, respectively). Afterwards, we used the getMethylationStats() command to obtain the percentage of methylated cytosines per context. Visualization of the methylated regions was done by converting alignment BED files into bigWig format with the Wig/Bedgraph-to-bigWig converter v1.1.1 included in Galaxy (Goecks et al., 2010). BigWig where then uploaded into Trackster, the visual analysis environment embed in Galaxy (Goecks et al., 2010). To check for symbodinium contamination we aligned with Bismark to the two available genomes (*S. minutum* GCA_000507305.1_ASM50730v1_genomic.fna rom NCBI and *S. kawagutii* Symbiodinium_kawagutii.assembly.935Mb.fa from http://web.malab.cn/symka_new).

## Results and discussion

### Coral nuclei are isolated with less than 2% contamination of Symbiodinium

Observation under fluorescent microscope showed that coral nuclei were not degraded and that there was little cell debris. We counted the number of intact *Symbiodinium* (which have a red fluorescence at 350 nm because of their chlorophyll content) and intact coral nuclei (with blue fluorescence caused by Hoechst 33342 binding to DNA). We observed the result of 3 different nuclei isolation and never found more than 2 intact *Symbiodinium* for 98 coral nuclei (2%). The same result was observed when trying with *Aiptasia sp.*. As expected, in menthol-bleached coral, no *Symbiodinium* were found. The low amount of contamination with *Symbodinium* was confirmed by WGBS: less than 0.004% of the total reads aligned to the *S. minutum* and *S. kawagutii* genomes.

### Mosaic DNA methylation occurs principally in CpG context in P. acuta

An average of 56.4 % (77,509,870 mapped read pairs out of 137,505,123 total read pairs) of the reads obtained for our three WGBS samples mapped on *P. acuta* reference genome. This may look like a small percentage of alignment, but this is classically observed with bisulfite sequencing (Gavery & Roberts 2010). Methylation occurs principally in CpG dinucleotide context, with 3.92±0.47 % (mean±SD, n=3) of them being methylated. 2.7% of CpG positions are highly methylated (≥70%), and about 81,000 positions (2% of analysed positions) are conserved *i.e.* in each of 3 replicates CpG sites are ≥70% methylated. In addition, 0.20 % of the CHG and 0.20 % of the CHH are methylated. Of the CHG and CHH positions less than 0.002% are methylated to ≥70% and no highly methylated conserved positions exist, *i.e.* CHH/CHG methylation is weak and dispersed. Total 5mC content of the genome is therefore roughly 0.5%. Methylation is distributed in a mosaic fashion, with large methylated intragenic regions (principally in exons) interspersed with large unmethylated regions. After visual inspection of the methylation profiles, we tentatively assigned a cut-off for highly methylated CpG to ≥80%. Highly methylated regions were defined as those with at least 5 highly methylated CpG separated by a maximum distance of 2kb between each of those CpG. Using these criteria, we identified 7,528 ± 146 highly methylated regions (**Figure 1**). The total length of these regions is 36,775 ± 1,436 kb corresponding to 14% of the genome. 18,399 (29%) of a total of 63,181 transcripts (Vidal-Dupiol et al. in preparation) correspond (at least partially) to these highly-methylated regions, indicating that highly methylated regions are enriched for transcribed sequences.

**Figure 1:**
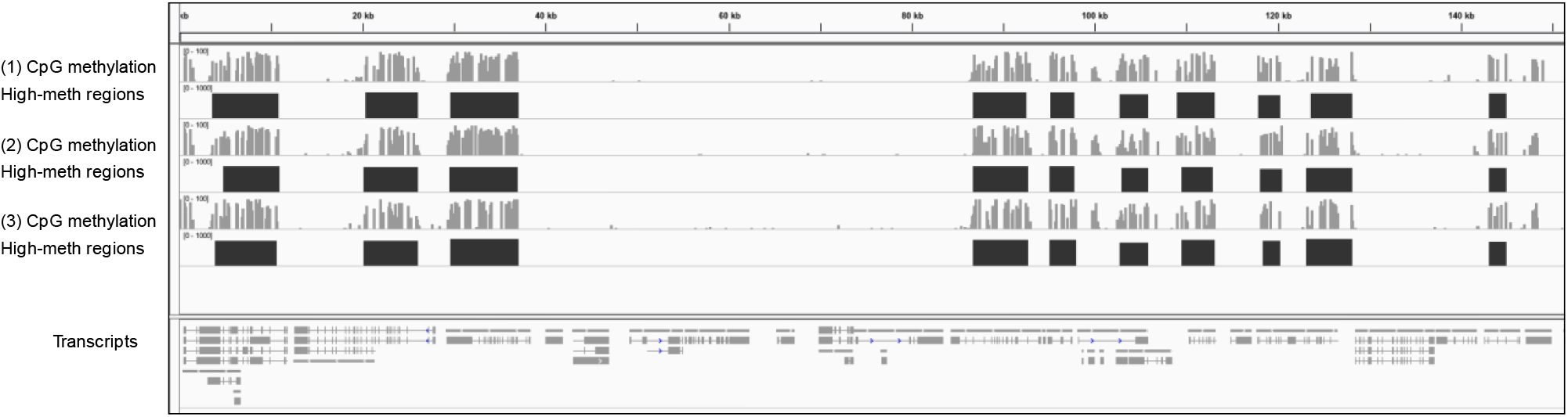
Example of methylation profiles deduced from BS-Seq in three replicates. Shown is scaffold150312_cov103 pos. 1-258,806 bp. Light grey profiles represent individual CpGs. Dark grey blocks represent regions that fulfill the criteria of highly methylated regions (max distance between CpG with at least 80% methylation is 2 kb, at least 5 CpGs present). Scale is 0 to max. For technical reasons max is 100% for CpG and a score of 1000 for regions (equivalent to 100%). Lower panel indicate transcripts.

### P. acuta possesses histones, but display an unusual chromatin structure shared with E. divisa and A. manjano

Transcripts for the four histones composing nucleosomes were found in *P. acuta* transcriptome and were aligned to other species to see similarity in amino acid sequences. *P. acuta* H3 and H4, have 94% and 98% identity in their amino acid sequences with *M. musculus*. Other two histones inside nucleosomes, H2a and H2b, have respectively 92% and 87% of identity with *M. musculus*. Histone tails for H3 and H4, where chemical modification occurs, were always 100% identical between mice and *P. acuta*. Sequence identity similarities were high for the four histones when compared those of the blood fluke *S. mansoni*, and similarities increased when sequence were confronted with more phylogenetically related organisms (*H. vulgaris* and *N. vectensis*). When compared to *A. digitifera*, another hermatypic scleractinian coral, 100% identity was found for *P. acuta* H2B, H3 and H4 protein sequences, and 99% with H2A.

We then performed western blots to determine if commercial antibodies targeting histone modifications could be used on *P. acuta*. We tested hamster brain (positive control), free *Symbiodinium* (negative control, as dinoflagellata contain no histone (Rizzo, 2003)), bleached and healthy corals. Results of the western blots are shown on **Figure 2.**

**Figure 2:**
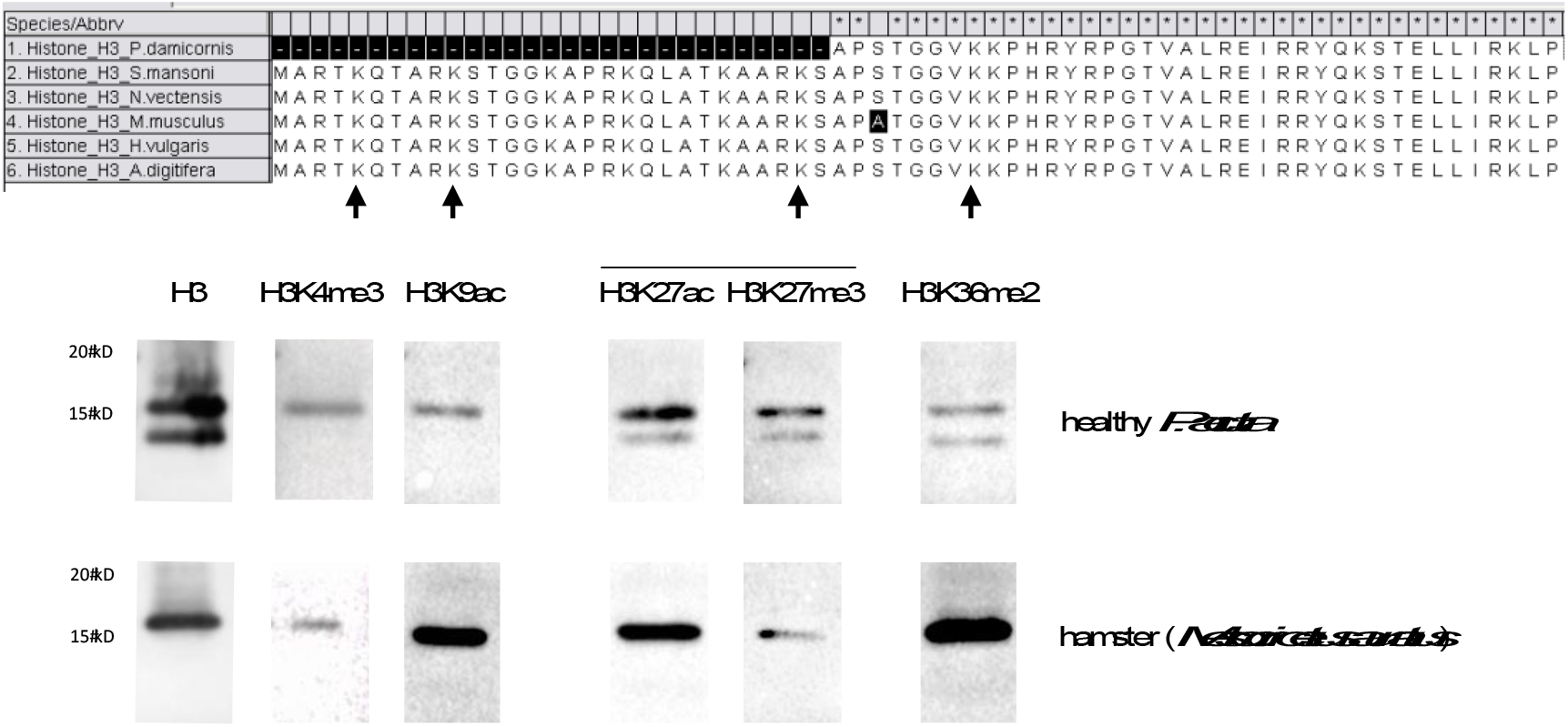
On top amino acid alignment of histone H3, positions 4, 9, 27 and 36 marked by arrow. H3 sequence of *P. acuta* predicted from genomic data. Black boxes indicate alignment differences and strockes missing data. Western blots using antibodies targeting histone H3, trimethylation of H3 lysine 4 and 27 of histone H3 (H3K4me3, H3K27me3), acetylation of H3 K9 and K27 (H3K9ac, H3K27ac), and dimethylation of lysine 36 of histone H3 (H3K36me2, right). Hamster *Mesocricetus auratus* as control.

Results for the three antibodies (anti-H3, anti-H3K27me3, anti-H3K36me2) were consistent between organisms. In all three cases, no signal was detected for *Symbiodinium,* and a signal at the expected size (approximately 15 kDa) for histone H3 was observed in hamster. In healthy corals, a signal was observed, but at 12 kDa. Bleached coral displayed both the 12 kDa band and the expected 15kDa band. The difference of about 3 kD corresponds roughly to a difference of 25 amino acid residues in size.

From these results, we suspected that there could be a different chromatin structure between bleached and healthy *P. acuta*, and we carried out a chromatin digestion with micrococcal nuclease (*M*Nase). *M*Nase digests DNA between nucleosomes, which creates DNA fragments with sizes that are multiples of approximately 150 bp (length of DNA wrapped around a nucleosome). When separated on a gel, it looks like a ladder. After prolonged digestion, only 150 bp fragments, corresponding to a mono-nucleosomal digestion, would be detectable on an agarose gel. A very good example of this can be seen in (Hewish and Burgoyne, 1973). We did not observe this type of result for neither healthy nor bleached *P. acuta*. Instead DNA is completely degraded with time, and no nucleosomal size fragments are seen (**Figure 3**).

**Figure 3:**
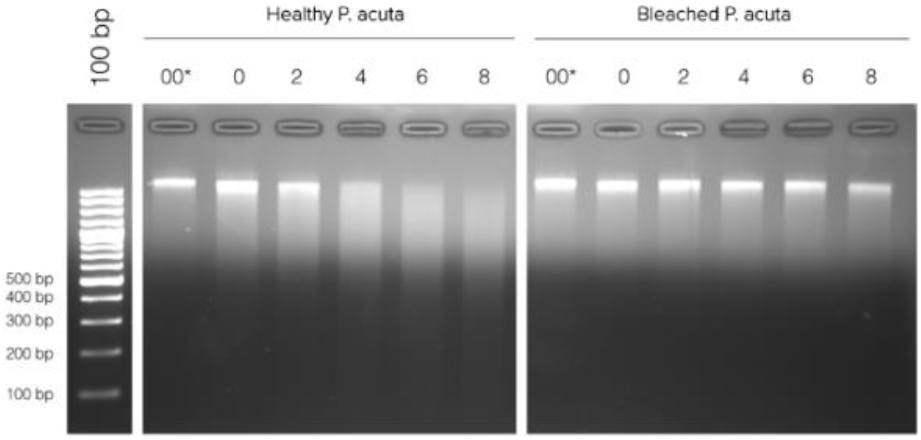
Micrococcal nuclease (*M*Nase) digestion profile of chromatin from healthy and bleached *Pocillopora acuta* in function of digestion time at 37°C. 00 corresponds to chromatin left at 37°C for 8 minutes, without the addition of *M*Nase, and 0 is addition of *M*Nase immediately followed by incubation on ice with EDTA (enzyme inhibitor).

To verify that it was not a problem with the chromatin extraction and digestion technique, we performed the same experiment on three other hermatypic corals *E. divisa*, *M. digitata* and *S*. *pistillata*, as well as two symbiotic anemones, *Aiptasia sp.* and *A. manjano.* As it can be seen on **Figure 4**, *E. divisa* and *A. manjano* present the same unusual digestion profile as *P. acuta*, while *S. pistillata*, *M. digitata* and *Aiptasia sp.* show the expected digestion profile, in the shape of a ladder and with an enrichment at 150 bp (corresponding to mono nucleosomal fragment). The ladder profile is not striking in *A. pallida* (probably because of an excess of starting biological material), but enrichment around 150 bp is clearly visible.

**Figure 4:**
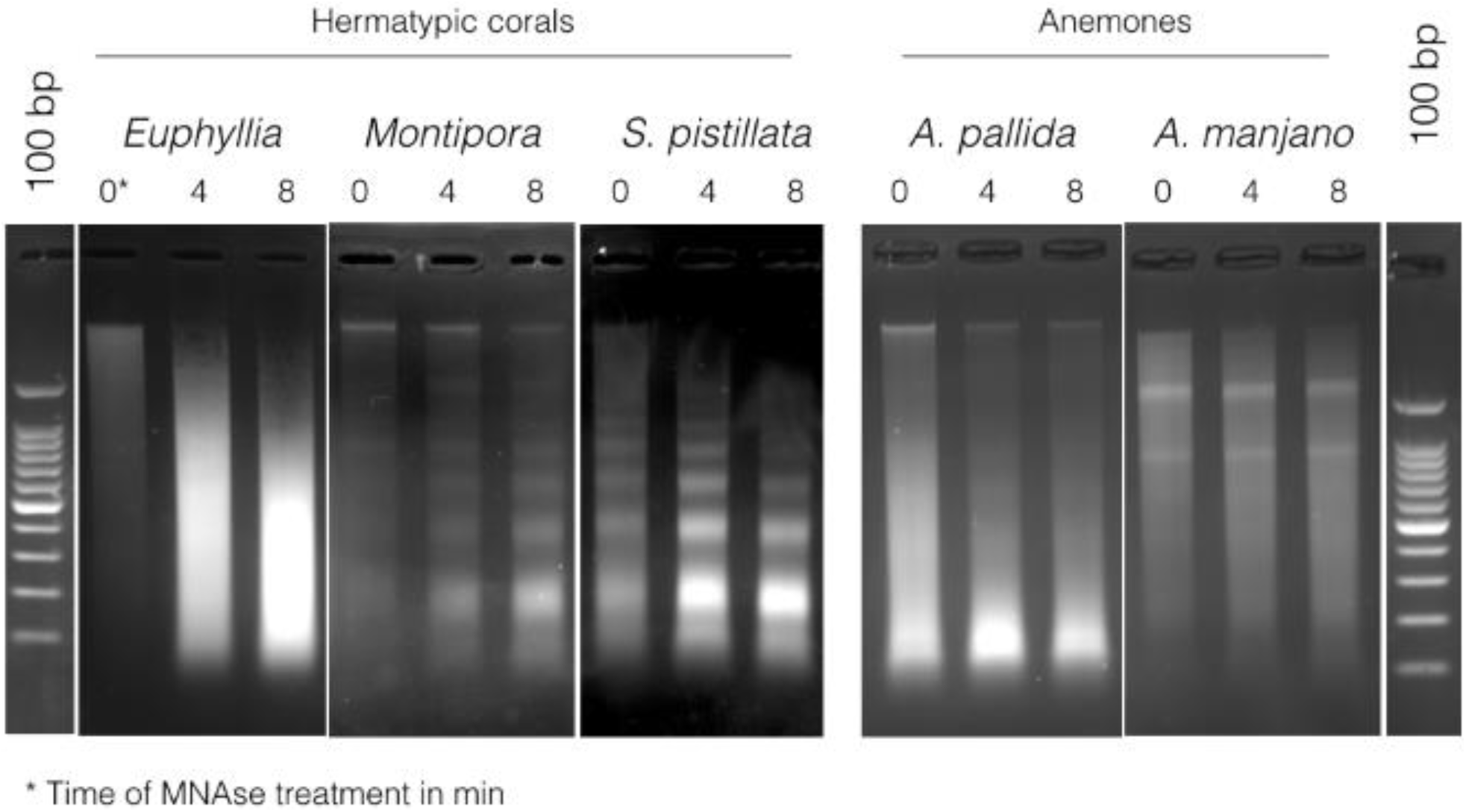
Micrococcal nuclease (*M*nase) digestion profiles for three hermatypic corals, *Euphyllia divisa*, *Montipora digitata.* and *Stylophora pistillata*, as well as for the two sea anemones *Aiptasia sp.* and *Anemonia manjano*.

## Conclusion

In this article, we describe (i) a straightforward method to separate *P. acuta* chromatin and DNA from *Symbiodinium* material, and (ii) provide the first description of a coral epigenome. We observed a “mosaic” DNA methylation pattern, with methylation-free regions interspersed with intragenic highly methylated ones (mostly on exons). Around 0.5% of the total cytosines are methylated, principally in the dinucleotide CpG. CHG and CHH methylation occurs nonetheless, but at lower frequency. These characteristics are fully concordant with other invertebrates in which DNA methylation is present and a methylome was described (Flores and Amdam, 2011; Fneich et al., 2013; Gavery and Roberts, 2010; Lyko et al., 2010; Suzuki and Bird, 2008; Xiang et al., 2010). Further analysis of the link between DNA methylation positions and functional elements of the genes has been done elsewhere (Vidal-Dupiol *et al*., in preparation).

We intended to use the isolated coral nuclei to perform chromatin immunoprecipitation (ChIP) on histone modifications. Alignments of protein sequences of histones H2A, H2B, H3 and H4, which compose nucleosomes, show that they are well conserved in *P. acuta*, with only little difference with mice. Interestingly, we did not observe the histone H3 band at the expected size (15 kDa) in healthy corals for any of the three antibodies we used (anti-H3, anti-H3K27me3, anti-H3K36me2). Instead, a single band of lower molecular weight (12 kDa ∆H3) is visible. Bleached corals are different, with both the expected H3 band at 15 kDa, and the truncated one at 12 kDa. The size difference of the bands suggests clipping of H3 tails at roughly 25 residues. Clipping after Ala21 in histone H3 was observed in *Saccharomyces cerevisiae*. Cleavage activity is induced under conditions of nutrient deprivation and sporulation (Santos-Rosa et al., 2008). H3 cleavage was also observed in vertebrates where it is cell type-dependent. Two clipping sites exist, one between Lys-23 and Ala-24 and the other between Lys-27 and Ser-28 (Mandal et al., 2013). Since we obtained a positive signal on Western blots for ∆H3 with anti-H3K27me3, cleavage occurs downstream of position 27 in *P. acuta*. Histone clipping is probably involved in the regulation of gene expression in a multitude of processes (Zhou et al., 2014). Surprisingly, ∆H3 is the major H3 isoform in healthy corals as opposed to bleached ones. As anticipated, no signal was detected in *Symbiodinium* extract, as they do not possess real histones (Rizzo, 2003). *Symbiodinium* live within coral cells, and their presence or absence may cause major cellular and nuclear reorganization as well as transcriptome remodelling. There is the possibility that *P. acuta* chromatin may be compacted in different fashion, with other histone forms or histone-like proteins, depending on its symbiotic state. Recently, Török et al. (2016) have shown that *Hydractinia echinata*, another Cnidaria, possesses 14 different histones (different from the canonical H1.1, H2A.1, H2B.1, H3.1 and H4.1 histones), including a novel H3.3.2 variant). These histones variants are expressed at different life stage or in diverse tissues to efficiently pack DNA. It is plausible that a similar diversity exists in *P. acuta* should be investigated. Further analyses are needed to confirm these hypotheses and to investigate if this phenomenon is also present in the other corals and anemone species used in this study. Surprisingly, treatment of the chromatin of *P. acuta* (healthy or bleached), *E. divisa* and *A. manjano* with *M*Nase did not produce nucleosomal fragments. *M*Nase cuts preferentially between nucleosomes because of steric hindrance. DNA is wrapped around nucleosomes for a 150 bp length, and incomplete digestion present a “ladder” profile on gels, with each bands being a multiple of 150 bp, while complete digestion shows a unique, intense band at 150 bp (Hewish and Burgoyne, 1973; Hörz and Altenburger, 1981; Keene and Elgin, 1981; Noll and Kornberg, 1977; Wu et al., 1979; Zaret, 2005). In these three species, DNA is completely digested by *M*Nase (**Figures 3** & **4**), similarly to what would happen in absence of nucleosomes (Hewish & Burgoyne 1973). On the other hand, *Aiptasia sp.*, *M. digitata* and *S. pistillata*, gave the expected digestion profile for *M*Nase. These results indicate that some species within the hexacorallia subclass have an unusual chromatin structure in which histones might be replaced by other proteins, or where nucleosomes are only loosely bound to DNA. However, these two digestion profiles are not coherent with the phylogeny of the species we used in this study (Daly et al., 2003; Romano and Palumbi, 1996), implying that it is not a feature specific of an order or a genus (**Figure 5**).

**Figure 5:**
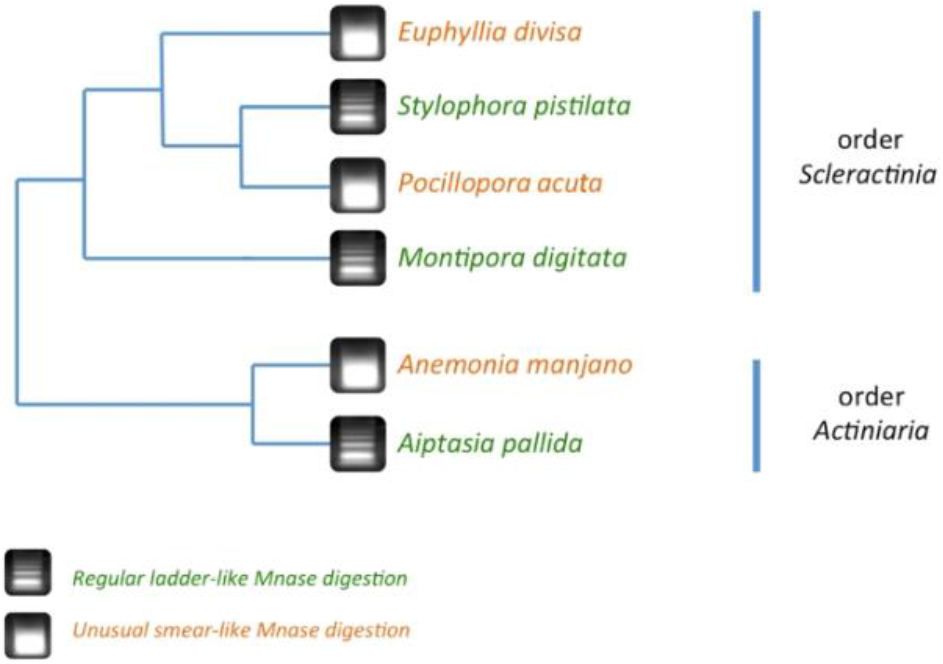
Simplified phylogeny of the coral and anemone species used in this study, based on (Daly et al., 2003; Romano and Palumbi, 1996) and associated with the results of the micrococcal nuclease (*M*nase) digestion. The digestions profiles do not follow the phylogeny, hinting that the unusual chromatin structure profile is not an ancestral conserved trait in the hexacorallia subclass.

At the moment, we cannot state if this unusual chromatin structure is an ancestral feature of the *Hexacorallia*, which was lost in some species, or if it is a new character that appeared at multiple times within the *Hexacorallia* subclass. More analyses are needed in order to see if this unusual chromatin structure is limited to some species in the *Scleractinia* and *Actinaria* orders, or if it is more widespread within *Hexacorallia* subclass or *Anthozoa* class.

## Author Contributions and Notes

JVD and CG designed research, DR, APP and KBR performed research, DR and CG analyzed the data, JVD, KBR, PR and PM collected material; and all authors wrote the paper.

The authors declare no conflict of interest.

## Acknowledgements

The authors are very grateful for the permission to use the aquarium facilities of the UMS2348 for the sample generation. This study was supported by the French “Agence Nationale de la Recherche” through the Program BIOADAPT (ADACNI ANR-12-ADAP-0016-03).

## Data Availability

Data for bisulfite sequencing are available as fastq files Sequence Read Archive (SRA) submission SUB1929299 at the NCBI SRA.

